# Derived motor neurons transplantation promotes nerve regeneration in a rodent model of a chronically denervated nerve through delayed adoption of a chronic denervation phenotype

**DOI:** 10.1101/2025.01.21.634102

**Authors:** Cashman R. Cashman, Ruifa Mi, Ahmet Hoke

**Affiliations:** MSTP/MD-PhD Program; Department of Neuroscience; Department of Neurology, Johns Hopkins School of Medicine, Baltimore, MD, 21205, United States

**Author notes:** Corresponding author: MIND Neurology Research, Building 114 Room 2000, 114 16^th^ St, Charlestown, MA, 02129, US. Department of Neurology, Massachusetts General Hospital, Boston, MA, 02114, US.

**Keywords:** Regeneration, peripheral nerve, stem cell, transplantation, chronic denervation, synaptogenesis

## Abstract

After acute injury, the peripheral nervous system regenerates more efficiently than the central nervous system, but this advantage is not maintained with chronic injuries. In large animals like humans, the slow rate of axon growth and the long distance between the central nervous system and end organ result in the adoption of a chronically denervated nerve phenotype even in the setting of acute injury: distal areas of the nerve become unable to support regenerating axons, even as the proximal aspect of the acute injury may be appropriately supportive of regeneration. We hypothesized that motor neuron transplantation into the denervated distal stump would maintain the regenerative capacity by effectively reducing the time and distance necessary for regeneration through relay formation as well as by supporting the host Schwann cells responsible for guiding regenerating axons. Using an *in vitro* assays, we showed the feasibility of relay formation by demonstrating ventral horn to derived-motor neuron synaptogenesis as identified by calcium imaging. Pilot studies of transplantation of embryonic stem cell-derived mouse motor neurons into chronically denervated rat nerves demonstrated improved regenerative capacity in the chronically denervated nerve following a delayed repair paradigm. While host to graft synaptogenesis was not observed, the Schwann cells in the transplanted nerve stump were maintained in a pro-regenerative state despite chronic denervation. These pilot data suggest cell transplantation can delay the adoption of a chronically denervated phenotype primarily by maintaining Schwann cells in a pro-regenerative state.

## Introduction

While regeneration of the peripheral nervous system is often considered to be robust, if a nerve becomes chronically denervated, recovery is severely diminished due to loss of Schwann cells. In a model of regeneration through a chronically denervated nerve, whereby the denervated distal stump of a nerve was cross sutured to a freshly transected proximal stump, a denervation of just eight weeks corresponded to a 40% reduction in the number of regenerated motor neurons, while a twelve week denervation lead to a nearly 80% reduction (Sulaiman and Gordon, 2000). The conversion from a pro-regenerative non-regenerative environment is in part due to loss of structural support through Schwann cell death and fibrosis (Weinberg and Spencer, 1978) as well as reduction of trophic factors (Höke et al., 2002). The loss of these regenerative cues ultimately leads to regenerative failure.

To support the regenerative capacity of a chronically denervated nerve, we propose direct transplantation of neurons into the distal nerve stump. The implanted cells may promote regeneration by serving as postsynaptic targets for regenerating host fibers to form a relay to the end organ (and thereby reduce time to regeneration) or preventing host Schwann cell degeneration. Previous transplantation experiments have suggested relay formation *in vivo* is possible. Transplantation of rat neural stem cells into the lesioned spinal cord of a rat showed electrophysiological evidence of relay formation between descending axons from the host and differentiating neurons from the graft (Lu et al., 2012). More recently, induced pluripotent stem cell derived neurosphere transplantation in a rodent model of spinal cord injury lead to activity facilitated synaptic integration into the rodent spinal cord with improved locomotor function (Kawai et al., 2021). It should be noted that in both of these experiments, host fibers were composed of upper cortical or brain stem axons, not lower motor neurons. The sufficiency for the peripheral nerve to support synaptogenesis, and the ability of lower motor neurons to synapse on other neurons has thus not been investigated. Additionally, transplanted neurons may support the endogenous Schwann cells such that they, in turn, do not atrophy and are capable of aiding endogenous neuron regeneration. The two mechanisms of relay formation and host support are not mutually exclusive.

To test the hypothesis that transplantation of motor neurons may maintain host regenerative capacity in chronic denervation, motor neurons derived from mouse embryonic stem cells were characterized and tested *in vitro* then ultimately transplanted into denervated nerves of rat hosts, with functional characterization following repair after critical delays of two to three months. Regeneration was assayed by imaging of the transplanted nerve and distal musculature, as well as functionally characterized by basic electrophysiological and gene expression studies.

## Materials and Methods

### Experimental Design

*In vitro* studies of purity were designed with multiple (3-5) differentiations to account for the natural variability of the differentiation process. Calcium imaging was performed with multiple cells from a single differentiation and multiple explants, since imaging was performed on a single cell level. Two- and three-month delays were intended to test the adoption of chronic denervation in the axon and Schwann cell, with three-month time point being more sensitive to Schwann cell death. *In vivo* studies included eight separate differentiations evenly distributes across all time points, with live cell, dead cell, and vehicle transplants randomly assigned to animals. All long-term *in vivo* studies were from the same passage mESCs as those studied in the characterization staining. While these studies were initially designed for a minimum of eight animals per time point, representing three or more different motor neuron differentiations, tumorigenesis reduced this to minimum of three. Expression studies were normalized to the contralateral nerve.

### Animals

All animal surgeries were conducted under protocols approved by the Johns Hopkins University Animal Care and Use Committee following the guidelines established by the National Institutes of Health and the American Association for the Accreditation of Laboratory Animal Care. All animals were housed under standard conditions. All rats were purchased from Charles River. Two cohorts of 40 and 41 ten-week-old Sprague-Dawley rats (Charles River; Wilmington, MA) were used for the main denervation/transplant/repair experiments, with 8-11 animals provided at each time point and four animals per treatment group. Any animal that developed a tumor was not included in the final analysis.

### Motor neuron differentiation, culture, and preparation for transplant

Motor neurons were differentiated from the HBG3 Hb9:GFP mouse embryonic stem cells (mESCs) based on previously published protocols (Wichterle et al., 2002; Wichterle and Peljto, 2008; Wichterle et al., 2009). Briefly, primary mouse embryonic fibroblasts (Chemicon, PMEF-N; Temecula, CA) were plated onto coated tissue culture plates at 160 cells/mm^2^ one day prior to mESCs (unless otherwise noted) with feeder media containing 10% ES FBS (Gibco, 10439-024; Grand Island, NY), 1x Glutamax (Gibco, 35050-061; equivalent to 2 mM glutamine) and 1x antibiotic/antimycotic (Gibco, 15240-062) in ES DMEM base media (Millipore, SLM-220-B). The mESCs were expanded on the feeders in mESC media containing 15% ES FBS, 1x antibiotic/antimycotic, 1x Glutamax, 1x nonessential amino acids (Millipore, TMS-001-C; Burlington, MA), 1x nucleosides (Millipore, ES-008-D), 100 μM β-mercaptoethanol (Sigma-Aldrich, M3148), and 5*10^5^ U rLIF (“ESGRO,” Millipore, ESG1106) in ES DMEM base media. mESCs were grown until the colonies were 60-70% confluent, at which point they were dissociated from the dish with 0.05% trypsin/EDTA (Gibco, 25300-054). The PMEFs were separated from mESCs by differential adhesion to a tissue culture plate whereby PMEFs bind more quickly than mESCs. Embryoid bodies were produced from the mESCs by resuspending in DFK10 media containing 10% Knockout serum replacement (Gibco, 10828-028), 1 x antibiotic/antimycotic, 0.5x Glutamax (equivalent to 1 mM glutamine), 0.15% glucose (Sigma-Aldrich, G7021; St. Louis, MO), 1x N2 supplement (Gibco, 17502), and 110 μM β-mercaptoethanol in a 50:50 base media solution of ES DMEM and F12 (Gibco, 11765) for two full days, with fresh media each day. After embryoid body (EB) growth, motor neuron differentiation was performed by culturing the cells in DFK10 media supplemented with 1 μM retinoic acid (Sigma-Aldrich, R2625) and 2 μM of the sonic hedgehog agonist purmorphamine (Millipore, 540220). Differentiation media was changed daily for five total days of motor neuron induction.

Following motor neuron differentiation, the cells were gently dissociated into a single cell suspension. Prior to dissociation, cells were treated with 50 μM ZVAD (Millipore, 627610) for 30 minutes at 37°C. EB/motor neurons were washed with PBS supplemented with 50 μM ZVAD then treated with cold 0.05% trypsin/EDTA until the EBs began to aggregate (about 5 minutes). The reaction was stopped with heat-inactivated horse serum (Sigma-Aldrich, H1138) then triturated in a 1:1:2 solution of ES DMEM:F12:Neurobasal (Gibco, 21103-049) with 25 μg/mL DNAse (Sigma-Aldrich, DN25) and 50 μM ZVAD. The cell solution was then passed through a 70 μM cell strainer (Corning, 431751) and pelleted.

Cells were resuspended in motor neuron media containing 5% horse serum (Gibco, 26050-070), 1x N2 supplement, 1x B27 supplement (Gibco, 17504-044), 1x antibiotic/antimycotic, 0.775x Glutamax (equivalent to 1.55 mM glutamine), in a 1:1:2 base media mix of ES DMEM:F12:Neurobasal media supplemented with 10 ng/mL BDNF (Life technologies, PHC7074; Carlsbad, CA), 10 ng/mL GDNF (R+D Systems, 212-GD-010/CF; Minneapolis, MN), 10 ng/mL CNTF (R+D Systems, 257-NT-010/CF), 2 ng/mL HGF (Sigma-Aldrich, H9661), 10 ng/mL IGF-1 (R+D Systems, 291-G1-200), 20 ng/mL basic FGF (Life technologies, PHG0024), 20 ng/mL acidic FGF (Sigma-Aldrich, F5542), 20 ng/mL EGF (Sigma-Aldrich, E1257 or Peprotech 315-09), 3 ng/mL PDGF (Peprotech, 100-13A; Cranbury, NJ), 50 ng/mL NT-3 (Peprotech, 450-03), 50 μM MDL28170 (Sigma-Aldrich, M6690), 50 μM ZVAD. Cells viability and concentration was determined with Trypan Blue staining (Corning, 25-900-CI) and a hemocytometer.

For transplant, cells were prepared at 10^5^ live cells/μL in fully supplemented motor neuron media then stored on ice or heated for 15 minutes at 100°C for heat-killed control cells. Cells for culture were plated on laminin-coated glass coverslips with or without glia, as indicated, at a density of 625 or 10^3^ cells/mm^2^ (lower density for general imaging, higher density for synaptogenesis experiments) and grown in motor neuron media for one day, then in media identical to growth media without ZVAD. Half media changes were made every 2-3 days.

### Ventral horn explant co-culture

Spinal cord explants were prepared from P8 rat or mouse pups. The lumbar spinal cord was isolated and then dissected in Hanks media (Gibco, 24020-117) with 143 mM glucose (Sigma-Aldrich, G7528) to remove roots and dura. After dissection, the ventral horn was separated from the entire cord along the central canal. After sectioning into 350 μm sections, the ventral horn tissue slices were manually cut into 6-8 pieces. These ventral horn sections were then transferred to a laminin coated Millicell Insert (Millipore, PIC0403050) and cultured with “Explant Media” containing 250 mM HEPES (Fischer, BP310-100; Pittsburgh, PA), 35.75 mM glucose, and 1x antibiotic/antimycotic (Gibco, 15240-062) in 50% MEM (Gibco, 11575-032), 25% Hanks media with 25% horse serum (Gibco, 26050-070) base media at pH 7.41 supplemented with GDNF to 10 ng/mL (R+D Systems, 212-GD-010/CF). All images of explants were of live samples with no fixation or staining.

### Immunostaining

Cells were fixed on coverslips with 4% paraformaldehyde (Electron microscopy services, 15714-S) in phosphate buffered saline (PBS) for 20 minutes at room temperature, rinsed three times with PBS, then stored at 4°C until staining.

Fixed tissue sections were prepared as either floating sections (muscle for neuromuscular junction staining, nerve for graft imaging) or mounted on gelatin coated glass slides (nerve for macrophage staining or muscle for myofiber diameter staining). To this end, following collection, nerve tissues and muscle tissues were fixed overnight at 4°C (nerve) or 20 minutes at room temperature (muscle) in 4% paraformaldehyde in PBS. After fixation, samples were transferred to 15% sucrose (Sigma-Aldrich, S5015) in PBS for 24 hours at 4°C, then a 30% sucrose solution for an additional day at 4C. Tissue samples were embedded in OCT cryoprotectant (TissueTek, 4583; Gardena, CA) and frozen at −80 for at least one day. Samples were then sectioned on a cryostat (Microm HM 505E Thermo Scientific; Walldorf, Germany) at −20°C at 30-50 μm (floating sections) or 10-12 μm (slide mounted sections). All floating sections were transferred to PBS inside of Netwell Inserts (Corning, 3479; Glendale, AZ) for staining. Glass mounted tissue sections represented consecutive sections. Samples were stored at 4°C until use.

Fixed cell or sectioned tissue samples were washed three times in PBS then permeabilized for 10-15 minutes with 0.1% Triton X-100 (Sigma-Aldrich, X-100). The samples were washed again and blocked for 30-60 minutes at room temperature in a buffer containing 5 or 10% normal goat or donkey serum (Cell Signaling Technology, 5425S; Danvers, MA; or Sigma, D9663, respectively), 0.5% Tween 20 (MP Biomedicals, MP1Tween201; Solon OH) in PBS. Primary antibodies were diluted in blocking buffer and incubated with samples overnight at 4°C. The next day, samples were thoroughly washed with PBS. Secondary antibodies were then prepared in blocking buffer and incubated with samples for one hour at room temperature. All samples were then mounted with mounting solution containing the nuclear stain DAPI (Vector H-1200; Newark, CA) and sealed with nail polish.

For the five-week-post transplant tissue, neurons/axons were stained with primary antibodies rabbit α-PGP9.5 and chicken α-GFP, with secondary antibodies goat α-chicken, AF488 and goat α-rabbit, Texas Red. Eight-week-denervated nerve samples were stained with primary antibody mouse α-β-III-tubulin and secondary goat α-mouse, Texas Red. Muscle samples were stained with primary antibody mouse α-β-III-tubulin and secondary labels horse α-mouse, FITC and α-bungarotoxin, AF594.

For complete nerve staining, fixed tissues were cleared following the CUBIC protocol (Tainaka et al., 2014). Briefly, whole, fixed tissue was placed in a “CUBIC-1” solution of 25/25/15% solution (by weight) of urea (Fisher, U15-500), *N,N’,N’*-tetrakis(2-hydroxypropyl) ethylenediamine (Sigma, 122262), and Triton X-100 (Sigma, T9284) at 37C for 3 days. The sample was then washed about four times for one hour each with PBS. Samples were then stained with primary antibodies rabbit α-PGP9.5 and chicken α-GFP, with secondary antibodies goat α-chicken, AF488 and goat α-rabbit, Texas Red in a solution of 0.1% Triton-X100 in PBS. Antibody incubation was for about 3 days at room temperature, with four one-hour washes between the primary and secondary. Stained samples were then incubated with “CUBIC-2” solution of 50/25/10/0.1% solution of sucrose (Fisher, S5500), urea, 2,2ʹ,2ʹ’-nitrilotriethanol (Sigma, 101447689), and Triton X100 for about three days at 4C in the dark.

For synapse staining, motor neurons were cultured from one to seven days, as indicated, and stained. More specifically, ChAT staining used primary antibody goat α-ChAT and secondary donkey α-goat, AF555 in 5% normal donkey serum blocking buffer. vGLUT staining was also performed in 5% normal donkey serum blocking buffer with primary antibodies goat α vGLUT1 and goat α-vGLUT 2 with secondary antibody donkey α-goat, AF555. PSD95 staining was performed in 10% normal goat serum blocking buffer with rabbit α-PSD and chicken α-GFP and secondary antibodies goat α-rabbit, Texas Red and goat α-chicken, AF488.

For qualitative images for comparison of GFP/ChAT to GFP/PSD95 and GFP/vGLUT, differentiated motor neurons after one day in vitro were stained for GFP with the rabbit α-GFP and ChAT goat α-ChAT primary antibodies with horse α-goat, FITC and donkey α-goat, AF555 (as detailed above). To avoid cross reactivity between the secondary horse α-goat, FITC and goat α-ChAT, GFP staining was completed prior to ChAT staining. Images were at 20x on an Observer.Z1 (Zeiss; Oberkochen, Germany) with LED excitation filtered to appropriate frequencies. Antibodies and dilutions detailed in Table 1.

**Table 1.** Antibodies.

| *Antibody name* | *Dilution* | *Manufacturer* | *Catalogue Number* |
| --- | --- | --- | --- |
| **Primary antibodies** |  |  |  |
| Goat α-ChAT | 1:100 | Millipore | AB144P |
| Chicken α-GFP | 1:500 | Aves (Davis, CA) | GFP-1020 |
| Chicken α-Nestin | 1:10000 | Aves | NES |
| Rabbit α-GFAP | 1:400 | Dako (Santa Clara, CA) | Z0334 |
| Rabbit α-GFP | 1:400 | Life technologies | A11122 |
| Rabbit α-Neurofilament | 1:400 | Millipore | AB1989 |
| Rabbit α-Olig2 | 1:500 | Millipore | AB9610 |
| Rabbit α-PGP9.5 | 1:1000 | BioRad (Hercules, CA) | 7863-0504 |
| Rabbit α-PSD95 | 1:200 | Cell signaling<br>technology | 3450S |
| Mouse α-Synaptophysin | 1:200 | Millipore | mab5258 |
| Mouse α-β-III-tubulin | 1:1000 | Promega (Madison, WI) | G7121 |
| Goat α-vGLUT1 | 1:400 | Abcam (Waltham, MA) | ab104899 |
| Goat α-vGLUT2 | 1:400 | Abcam | ab101760 |
| **Secondary Antibodies** |  |  |  |
| α-Bungarotoxin, AF594 | 1:1000 | Life technologies | B13423 |
| Goat α-Chicken, AF488 | 1:500 | Life technologies | A11039 |
| Goat α-Chicken, AF594 | 1:500 | Life technologies | A11042 |
| Donkey α-Goat, AF555 | 1:400 | Life technologies | 21432 |
| Horse α-Mouse, AMCA | 1:400 | Vector Labs | CI-2000 |
| Horse α-Mouse, Texas Red | 1:400 | Vector Labs | TI-2000 |
| Goat α-Rabbit, FITC | 1:400 | Vector Labs | FI-1000 |
| Goat α-Rabbit, Texas Red | 1:400 | Vector Labs | TI-1000 |

Quantitation of ChAT versus GFP^+^ population was performed from 10x Z-stack images of differentiated motor neurons on laminin, rat astrocyte, and rat Schwann cell substrates after one week *in vitro*. No GFP antibody was used for the quantitation to avoid bleed through of the green into the red channel.

Note that all secondary antibodies were also validated against no primary controls to assess background activity. Primary and secondary antibodies with sources and dilutions are below, Table 1.

### *In vitro* synaptogenesis and calcium imaging

Triple cultures were prepared for both synaptic agonist determination of synapse identify as well as synaptic antagonists following electrical stimulation. These were prepared slightly differently, as outlined below.

Agonist studies for synapse characterization were performed by bath application of three synaptic agonists and a positive control to a triple culture of ventral horn explant, cellular substrate (rat Schwann cells. Spinal cord explants were prepared as outlined above and cultured in “SC Explant Media +GDNF” (see above) until the addition of mESC-derived motor neurons at which point it was changed to motor neuron media without ZVAD, unless otherwise indicated. Mouse astrocytes were isolated from P1 mouse pups and cultured on poly-L-lysine/matrigel coated dishes for 1.5 weeks prior to initiation of glia/motor neuron co-culture experiments. Rat Schwann cells were passage 11 from an isolation from a P4 rat pup. Glia were plated at 250-500 cells/mm^2^ and allowed to expand for 1-7 days prior to addition of motor neurons. Motor neurons were differentiated and dissociated as described above. At the time of dissociation, they were resuspended in supplemented motor neuron media + ZVAD and 50 ng/mL GRO-1 (Peprotech, 300-11) then plated onto the glia at 10^3^ cells/mm^2^. The culture then grew for seven days with daily 50% media changes prior to calcium imaging.

Cells were loaded with Fura-2 by incubating the cells at 37°C for 30 mins with a 1 mM Fura-2AM (Life technologies, F1201) in a 2% bovine serum albumin (Sigma, A9647) solution in artificial cerebrospinal fluid (aCSF). aCSF was prepared as a solution of 140 mM NaCl (JT Baker, 3624-01), 4.8 mM KCl (Sigma-Aldrich, P3911), 1.0 mM MgCl_2_ (JT Baker 2444-01), 2.0 mM CaCl_2_ (Sigma-Aldrich, C3881), 40.0 mM D-glucose (Sigma-Aldrich, G7528), and 10.0 mM HEPES (Sigma-Aldrich, H4034-100G).

Synaptic agonists were prepared in aCSF as 50 μM glutamate (Sigma-Aldrich, G5667), 500 μM acetylcholine (Sigma-Aldrich, A2661), 100 μM serotonin (Sigma-Aldrich, H9523), and 10 mM aminopyridine (4AP; Sigma-Aldrich, 275875).

Cells were imaged at 20x on an Observer.Z1 (Zeiss) with a Lamda DG-4 light source (Sutter Instrument Company; Novato, CA) and physiological analysis performed by AxioVision (Zeiss). Calcium transients were acquired at 37°C. Exposure time following 340 and 380 nm excitation was adjusted such that the starting, baseline ratio of fluorescence was 1. Images with both 340 and 380 nm excitation were acquired every 1-5 seconds with the shorter interval during bath application. Agonists were perfused through the imaging chamber for bath application while images were acquired. Bath was maintained for two to five minutes then washed with aCSF for an additional two to five minutes (until fluorescence returned to baseline). Finally, 4AP was applied at 10 mM for one minute followed by a wash of 2-10 minutes (until baseline fluorescence returned).

Functional synaptogenesis was tested by electrical stimulation in the setting of bath application of synaptic blockade. Ventral horn explants were prepared as above and cultured for one week in explant media, after which primary rat Schwann cells were added at a density of 400 cells/mm^2^ and cultured in supplemented mouse motor neuron media for one day. Differentiated mouse motor neurons were plated at a density of 10^3^ cells/mm^2^ and cultured in supplemented motor neuron media with ZVAD and GRO-1. After one day in culture, the media was no longer supplemented with ZVAD. Stimulation and synaptic blockade were performed after one week of triple culture.

Synaptic antagonists were prepared in aCSF as a glutamate blockade containing 50 μM CPP (Tocris, 0173; Abington, England) and 50 μM NBQX (Tocris, 1044) as well as cholinergic blockade with 50 μM mecamylamine (Sigma-Aldrich, M9020). Electrical stimulation was performed with a handheld stimulator (ITO physiotherapy and rehabilitation, ES-130; Weymouth, MA) with 100 μS, bipolar 18V pulse applied at 80 Hz for 30 seconds. Maximum current output was 36 mA. The electrical stimulation was through two 30 or 36 gauge electroacupuncture needles (Seirin 020×30 or Electro-therapeutic devices HWA-TO, respectively; Shizuoka, Japan and Markham, Canada) positioned on either side of the ventral horn explant tissue (see Figure 2). Images were obtained at one second intervals. Wash and inhibitor baths were alternated. Inhibitor baths were applied for three to five minutes while wash was for a minimum of one minute. Electrical stimulation was performed two to three times in each bath. A final solution of 10mM 4AP was used, as previously, as a positive control and for normalization of the calcium transients.

Analysis was performed in AxioVision whereby GFP^+^ or GFP^-^ cell bodies were defined as a region of interest and the fluorescence measured over time. Additional analysis was performed in Microsoft Excel. Calcium transient traces (for agonist studies) were prepared by defining the first minute of fluorescence as baseline and subtracting the average value of this period from each individual time point, then dividing by the average baseline fluorescence (ΔF/F). All values were than normalized to the maximum response from 4-aminopyridine depolarization, to correct for signal attenuation from GFP (Bolsover et al., 2001). For quantitation of depolarization (for electrical stimulation and synaptic antagonist studies), baseline was defined as the minute prior to chemical or electrical stimulation, and the maximum response was defined as the maximum fluorescence until the sample returned to baseline. These responses were again corrected to 4-aminopyridine facilitated depolarization. The final value was an average of the response to all electric stimulations for a given condition. Reduction in depolarization evoked from electrical stimulus following synaptic blockade was determined by comparison to the response to a stimulation immediately prior to application of blockade.

Correlations were prepared as outlined in Statistical Methods.

### Imaging

Widefield epiflourescent, transmitted light, and live cell images of tissues or cells were obtained on an Olympus IX51 inverted microscope with a mercury bulb (Olympus, BH2-RFL-T3; Tokyo, Japan) and white light source (Olympus, JH40100). Fluorescent images were obtained with GFP, Texas Red, and DAPI filter sets.

Confocal images were obtained on a Zeiss LSM 510 META confocal microscope equipped with argon and helium/neon lasers for 543, 488, and 633 nm excitation. Cy5, rhodamine, and FITC filter sets were used for fluorophores of far-red, red, and green excitation/emission parameters, respectively.

For direct comparisons within a group, acquisition parameters of each channel were adjusted equally for all images.

Electron micrographs were obtained at 8000x on a Zeiss Libra 120 Plus Transmission Electron Microscope.

Image processing was performed with the Fiji software package (Schindelin et al., 2012; Schneider et al., 2012; Schindelin et al., 2015) to combine color channels. For direct comparisons within a group, digital gain of each channel was adjusted equally for all images. Z-stack projections were performed with the maximum intensity method. Co-localization of two channels was performed using the Coloc 2 plugin in Fiji (with region of interest defined around neurites of GFP^+^ cells).

### Denervation, transplantation, and repair

Experiments involving animals were approved by the Johns Hopkins University Animal Care and Use Committee and carried out according to relevant guidelines and regulations in a facility, which is accredited by the American Association of the Accreditation of Laboratory Animal Care. For all surgeries, anesthesia was induced with 2.0-2.3% isoflurane/oxygen (Piramal Healthcare; Mumbai, India). Animals were from seven to ten weeks of age at the time of transplant.

For denervation, the biceps femoris muscle was exposed following a 2.0-2.5 cm incision along the midline of the left upper hindlimb (from the sciatic notch to patella) after shaving and chemical depilation (Nair Church and Dwight; Ewing, NJ). The muscle’s insertion to the femur was transected and traction applied to display the sciatic nerve and its trifurcation into the common peroneal, sural, and tibial nerves. Connective tissue around the nerve was removed with spring scissors and 5-0 forceps (Fine Scientific Tools; Foster City, CA). Four ligatures were applied with 4-0 nylon sutures around the proximal most aspect of the tibial nerve. The nerve was then transected between the middle two ligatures. The proximal and distal stump were reflected in opposite directions. While the distal stump was loosely sutured to the adductor magnus with an extra loop from the penultimate distal ligature, the proximal stump was buried into the adjacent biceps femoris using two to three 10-0 polyamide or nylon sutures. Some animals (the four-week post-transplant engraftment analysis), denervation was simplified by double ligating the nerve with 10-0 polyamide suture, transecting the nerve between the ligations, and capping the distal stump to an electrospun polycaprolactone (80000 molecular weight, Sigma-Aldrich 440744) cuff sealed at the proximal end of the tube. The tube was 2-3 mm in diameter and 7 mm in length. The nerve stump/cap was then sutured to an adjacent muscle with 6-0 nylon sutures. After denervation, the muscle and skin were returned to their anatomical position and the incision closed with surgical staples. Staples were removed in all animals within four weeks.

For transplantation, the denervated tibial nerve stump was exposed as for the denervation surgery, one week after denervation. Two 1 μL injections of a 10^5^ live or dead cells/μL solution in motor neuron media (with 50 μM ZVAD) were made with a 5 μL syringe equipped with a 33-gauge needle (Hamilton, 65460-02; Reno, NV) into the distal stump of the tibial nerve one week after denervation. Vehicle transplants corresponded to two 1 uL injections of motor neuron media (with ZVAD) only. To facilitate insertion of the needle, a small glass micropipette was used to puncture the epineurium. Injections were placed two and seven millimeters distal to the distal most ligature with needle insertion into the endoneurial/perineural space. Following injection, the entry site was held closed with forceps for 30 seconds before needle extraction. After transplant, the surgical site was closed as above.

For repair, two or three months after transplant, the denervated distal tibial nerve stump was repaired by cross suture to a freshly transected common peroneal nerve stump. The nerves were exposed as above, and scar tissue manually dissected. The tibial nerve stump was cut from its ligatures and the proximal most millimeter trimmed. The common peroneal was then transected at the level of the distal tibial nerve stump. Three to four 10-0 nylon or polyamide sutures were used to connect the proximal common peroneal nerve stump to the distal tibial nerve stump end-to-end. The surgical site was closed, as above.

### Immunosuppression

Animals were immunosuppressed with subcutaneous injections of a cyclosporine solution (“Sandimmune,” Novartis; Basal, Switzerland) at a dose of 10 mg/kg. The drug was diluted in normal saline (Quality Biological, 114-055-101) to 10 mg/mL to facilitate injection. In the long-term denervation/transplant/repair animals this dose was reduced to 5 mg/kg following repair to reduce toxicity. Immunosuppression was initiated three days prior to transplant, and animal mass obtained once per week, adjusting dosing each week as needed.

### Compound muscle action potentials

Compound muscle action potentials (CMAPs) were obtained with either insertion of needles subcutaneously or direct nerve stimulation. In the former case, stimulating electrodes were placed at the sciatic notch, while in the latter, the electrodes were bent into a semicircular cuff and rested on the nerve to stimulate. The distance between the electrodes in the cuff was about two millimeters, while insertion points for stimulation at the sciatic notch were five to eight millimeters apart. Sciatic notch stimulation was only used for qualitative assessment of recovery and relay formation, while all quantitation was performed with CMAPs elicited by direct nerve stimulation at the graft site in the tibial nerve (for studies of engraftment before repair) or immediately proximal (within five millimeters) to the repair site on the common peroneal nerve (for studies of regeneration). CMAP recordings were obtained from the intrinsic foot muscle with insertion of the recording electrodes perpendicular to the long axis of the paw, under the distal most and middle sets of foot pads. A grounding electrode was inserted under the skin of the tail.

CMAPs were elicited with a stimulation of 1.00 ms delay, 0.10 ms duration with amplitude voltage ramped from 0.25 to 10v. Individual CMAPs were recorded at each step of the ramp. Data collection and analysis were performed in LabChart (AD Instruments; Dunedin, New Zealand). Peak-trough amplitude was calculated from the particular CMAP from the minimum plateau voltage (that is, the minimum voltage at which the CMAP ceased to increase in amplitude despite greater stimulation voltage). The high degree of scarring facilitated volume conduction, so higher stimulation amplitudes were not useful. Final CMAP amplitudes were normalized to values at time of repair for given treatment group.

For the longitudinal study of the emergence of the second peak, stimulation was at the sciatic notch at all but the eight weeks after repair time point. Quantitation and analysis were unchanged from the other samples.

### Quantitative qPCR

Using a QuantiTect Reverse Transcription kit (Qiagen, 205311; Venlo, Netherlands), cDNA was produced from 1 μg of total RNA extracted with Trizol (Life Sciences, 15596018) from tissue samples or dissociated cells. Tissue samples were homogenized with 1.6 mm steel beads (Next Advance, SSB16-RNA), and cell pellets were homogenized with trituration. Amplification was performed with QuantiScript SYBER Green PCR kit (Qiagen, 204143) as per manufacturer instructions, with each primer at a final concentration of 200 nM (400 nM for the pair). Samples were run in duplicate or triplicate on a StepOnePlus (Applied Biosystems; Waltham, MA) thermocycler and amplified as follows: 95°C for a 10 minute incubation, 40-50 cycles of 95°C for 15 seconds, 60°C for 1 minute. qPCR primers obtained through Sigma-Aldrich KiCqStart (KSP112012) with cJun, ErbB3, p75, and S100β and verified to not produce an amplicon with a negative control template (no reverse transcriptase), sequences summarized below (Table 2)

**Table 2:**
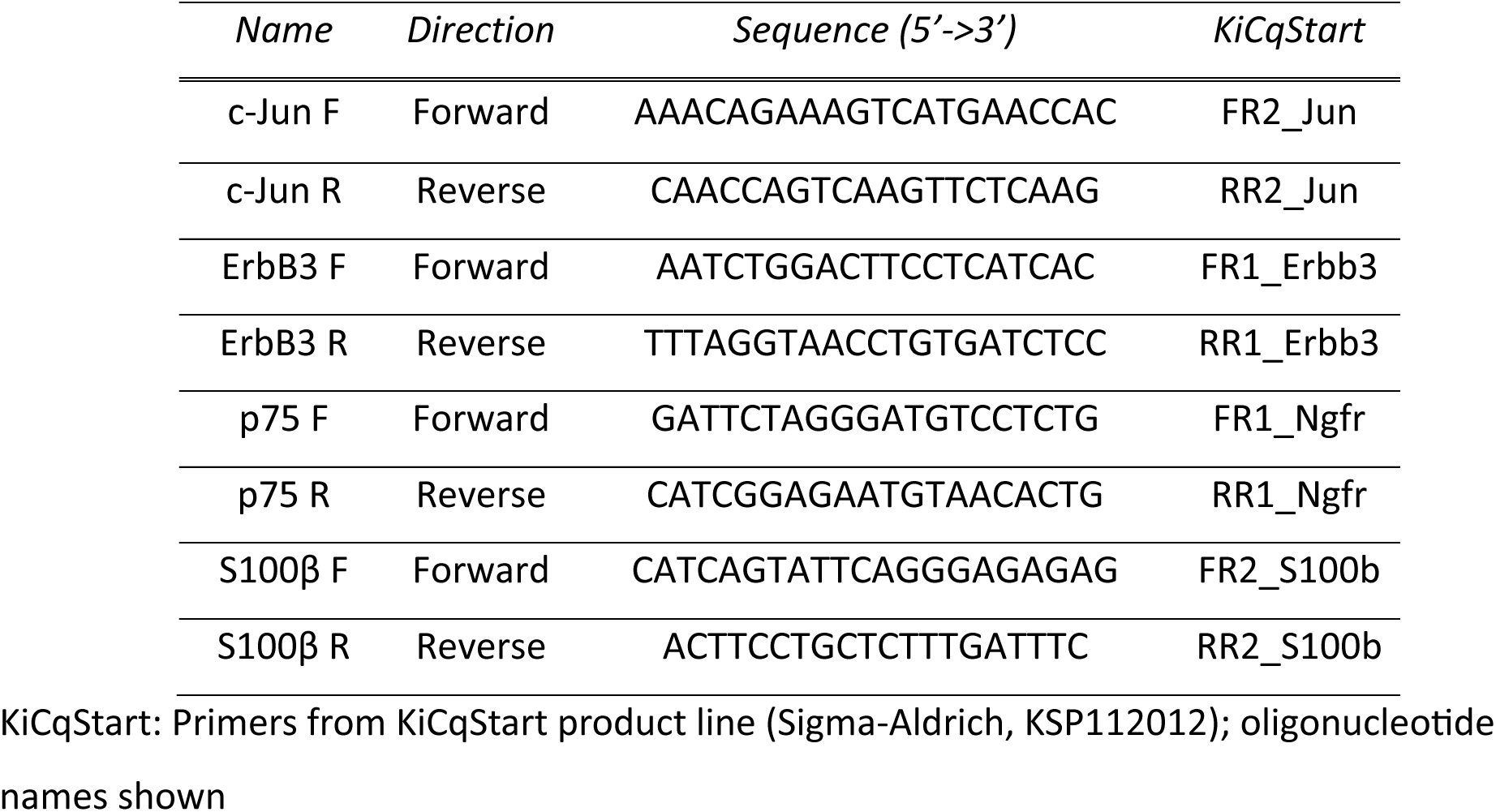
qPCR primer sequences.

Analysis was performed with the ΔΔCt method (Livak and Schmittgen, 2001) with genes of interest normalized to S100β and enrichment calculated relative to the intact side. The enrichment of the Schwann cell makers was then calculated relative to the dead cell transplant values for each gene of interest at each time point.

### Statistical tests

All animals that developed tumors were eliminated from analysis. After data collection but prior to analysis, data was tested for outliers using Grubb’s test with α=0.05 (GraphPad, Boston, MA). Outliers were removed prior to final statistical testing.

Multiple comparisons were tested by ANOVA using the R Studio software package version 2.15.2 (RStudio, 2015). If P<0.05 by AVOVA, Tukey’s post-hoc analysis was performed. All Student’s t-tests were performed in Microsoft Excel with homoscedastic (equal variance) or heteroscedastic (unequal variance) unpaired, two-tailed tests based upon the results of a F-test (null hypothesis accepted suggests equal variance), also performed in Excel. 95% confidence were prepared in Microsoft Excel using a Student’s t-distribution.

For correlation analysis of triple culture in different synaptic baths, Microsoft Excel was used to determine a line of best fit as well as Pearson’s correlation, with a manual Bonferroni correction for six Pearson’s correlations.

## Results

Motor neurons were differentiated from mouse embryonic stem cells (mESCs) with a motor neuron reporter Hb9:GFP using standard protocol of sonic hedgehog and retinoic acid induction (Wichterle and Peljto, 2008). The reporter enables rapid identification of motor neurons following differentiation and facilitates *in vitro* and *in vivo* imaging. Immunofluorescent staining was used to identify the cell type composition of four to seven separate differentiations from the same mESC passage number. Staining included anti-GFP staining to mark motor neurons (amplifying endogenous reporter), anti-beta-III tubulin for all neurons, anti-olig2 for oligodendrocytes, anti-GFAP for astrocytes, and anti-nestin for neural stem cells (Figure 1A). Nearly 15% of all cells were identified as motor neurons (GFP^+^/ß-3-tubulin^+^), and the majority of cells were neural stem cells (GFP^-^/Nestin^+^). A minority of cells were differentiated glia: oligodendrocytes (GFP^-^/Olig2^+^) and astrocytes (GFP^-^/GFAP^+^), Figure 1B. These data suggested the differentiated cells provided a reproducible and sufficient source of derived motor neurons.

**Figure 1:**
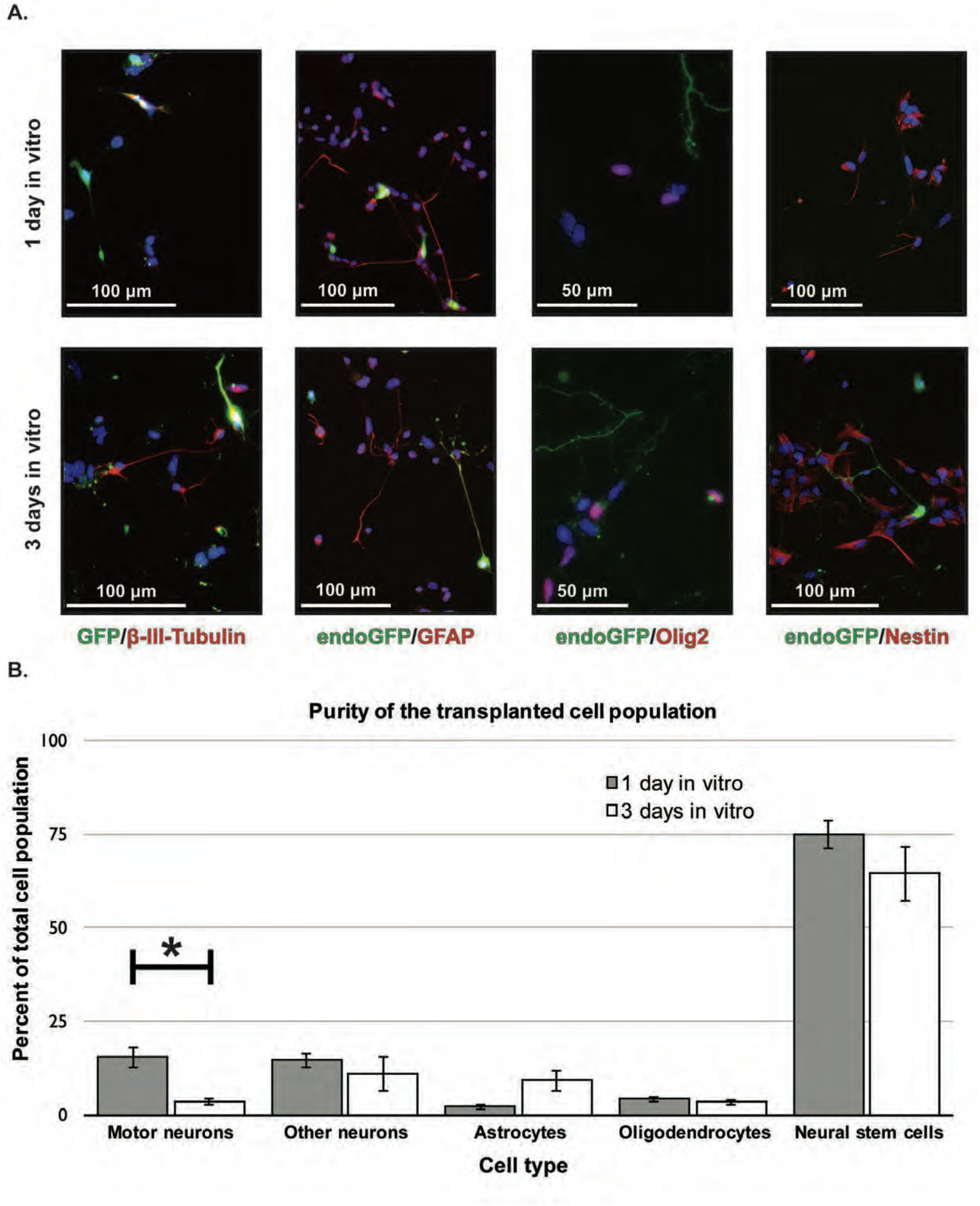
Characterization of the differentiated cell population A. Immunofluorescent images of motor neuron (α-GFP in green, left panel, 10x), neuron (β-III-Tubulin in red, first column, 10x), astrocyte (α-GFAP in red and endogenous GFP, second column, 10x), oligodendrocyte (α-olig2 in red and endogenous GFP, third column, 20x), and neural stem cells (α-nestin in red and endogenous GFP, fourth column, 10x) after 1 and 3 days *in vitro*. Images cropped to highlight cell morphology. Note for astrocytes, oligodendrocytes, and neural stem cells green is endogenous GFP without amplification by antibody staining (to avoid false positives due to secondary antibody background labeling). B. Quantification of cell purity versus all cells at 1 and 3 days *in vitro*. *=p<0.05 by Student’s two-tailed t-test with unequal variance. n=7 differentiations for motor neurons and neurons, n=4 differentiations for astrocytes, oligodendrocytes, and neural stem cells. All differentiations from same passage mESCs that were most used in this study.

### Synaptogenesis

The presence of appropriate synaptic infrastructure on the differentiated motor neurons was confirmed by immunofluorescent staining. When cultured on laminin, rat Schwann cells, or mouse astrocytes, the derived motor neurons (GFP+) expressed choline acetyl transferase (ChAT), essential for cholinergic transmission (2A), at 100%, over 75%, and 100% of motor neurons on Schwann cell, astrocyte, and laminin substrates, respectively (Figure 2B). ChAT^+^ but GFP^-^ cells are consistent with an interneuron identity. Derived motor neurons elaborated presynaptic (vGlut1/2) and postsynaptic (PSD95) structures necessary for glutamatergic transmission (Figure 2A). Pearson’s correlation of ChAT, vGlut1/2, or PSD95 co-localization with GFP (motor neuron) demonstrated high concordance between ChAT and GFP, with lower values for vGlut1/2 or PSD95 with GFP, consistent with the punctate staining of vGlut1/2 and PSD95 (Adler and Parmryd, 2010).

**Figure 2:**
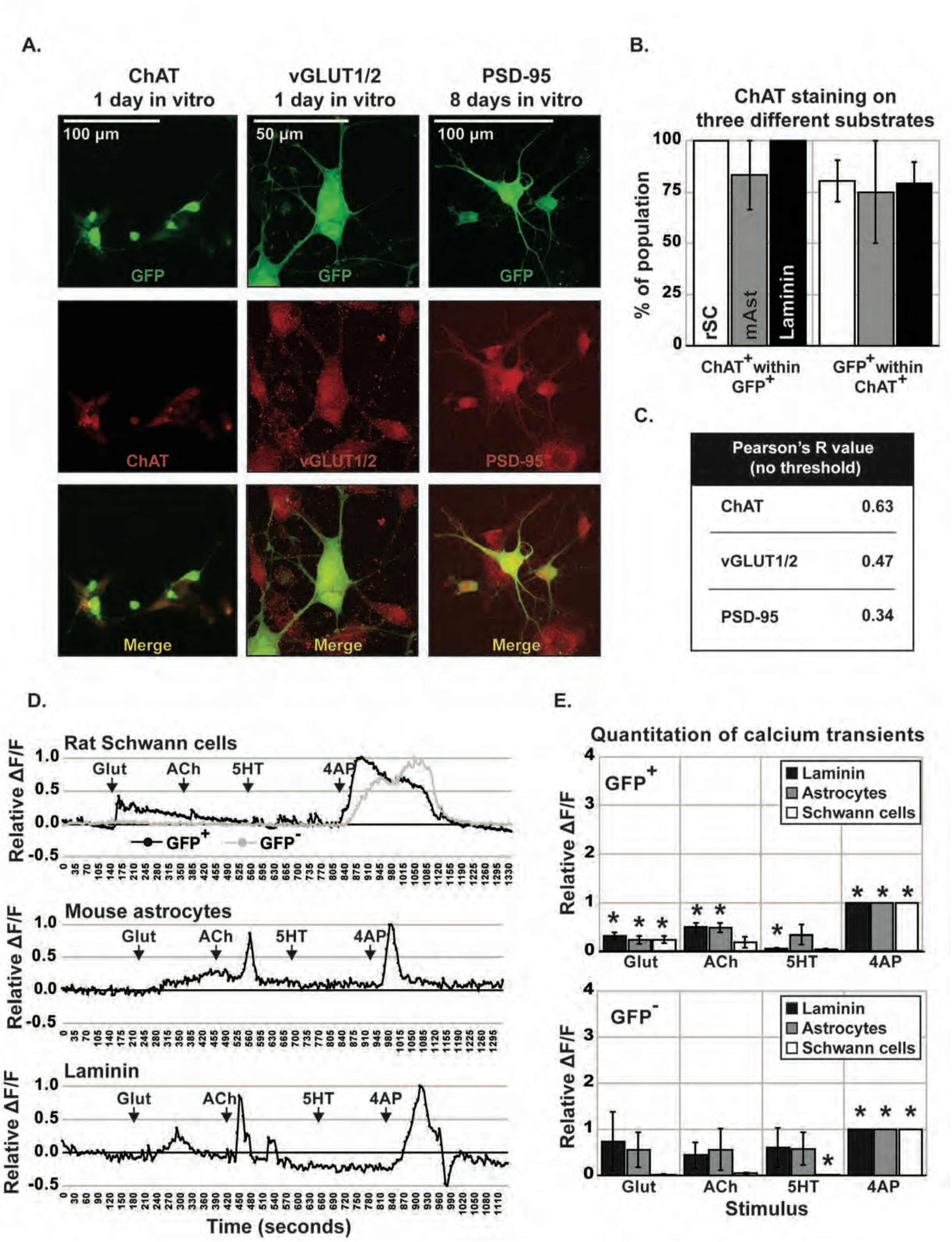
Characterization of glutamatergic and cholinergic synapses on differentiated motor neurons (**A**). Immunofluorescent confocal maximum intensity Z-projections of mESC-derived motor neurons at one or eight days *in vitro* on laminin with synaptic markers in red, as indicated. All samples had GFP amplified by α-GFP staining. Magnification as indicated. Note that ChAT staining is cropped to highlight two cells. (**B**). Quantification of ChAT and endogenous GFP double labeling on three different substrates after seven days *in vitro*. rSC: Rat Schwann cell, mAst: Mouse astrocyte. n=3 coverslips for all conditions. (**C**). R values of Pearson’s correlation of co-localization of red channel within green for the three merged images shown in A. Note that co-localization was performed with the Z-stack intact (where applicable) with regions of interest around the GFP^+^ cells and neurites only. (**D**). Calcium transients following neurotransmitter addition after motor neurons were cultured on three different substrates for seven days. Change in fluorescence ratio normalized to the maximum change following addition of 4-aminopyridine (10 mM). Top trace also shows the change in fluorescence ratio for a GFP^-^ cell. (**E**). Quantitation of maximum calcium response following neurotransmitter addition. GFP^+^ cell n=7, 5, and 7 cells on laminin, astrocytes, or Schwann cells, respectively. GFP^-^ cell n=3 cells on all three substrates. * is significantly different from 0 by 95% confidence interval with a Student’s t-distribution, α=0.05. Glut: glutamate, ACh: acetylcholine, 5HT: Serotonin, 4AP: 4-aminopyridine.

To determine if these synaptic structures were functional, cellular depolarization to glutamate and acetylcholine was tested through calcium imaging. Mouse embryonic stem cell-derived motor neurons were co-cultured with primary rat Schwann cells, primary mouse astrocytes, and a laminin-only substrate for seven days prior to calcium imaging, with the rat Schwann cells recapitulating future transplant experiments and mouse astrocytes being the natural support cell of the mESC-derived motor neurons. Calcium transients were visualized using the Fura2-AM calcium sensitive fluorescent dye. Responses of GFP^+^ or GFP^-^ cells to glutamate (50 μM), acetylcholine (500 μM), serotonin (100 μM), and 4-aminopyridine (10 mM) were measured on the three different substrates (representative traces, Figure 2D). On all three substrates, GFP^+^ cells responded to glutamate and, less consistently, to acetylcholine, as measured by normalized fluorescence difference. All cells depolarized to 4-aminopyridine (Figure 2D). A response of a GFP negative cell on rat Schwann cells is shown for comparison. Quantitation of calcium transients (normalized fluorescence difference before and after treatment) for all GFP^+^ cells showed significant (greater than zero by 95% confidence interval) responses above baseline to glutamate (Figure 2E, top). The response to acetylcholine was similar, reaching a significant difference from baseline in motor neurons on astrocytes and laminin. GFP negative cells had greater variability with a statistical response only to 4-aminopyridine (Figure 1E, bottom). While the response to serotonin (5-HT) was found to be statistically significant for GFP^+^ cells on laminin and GFP^-^ cells on Schwann cells, they were likely not biologically significant responses with mean values of 0.05 and 0.015, respectively. To determine if the synaptic machinery characterized above was sufficient for functional spinal cord to mESC-derived motor neuron synaptogenesis, a co-culture system was developed.

Rat spinal cord explant sections were co-cultured on rat Schwann cells with dissociated, differentiated motor neurons to determine if electrical depolarization of the explant could lead to chemical depolarization of the mouse motor neurons. The triple culture was started with a ventral horn explant, which was cultured under standard conditions for seven days. The subsequent addition of rat Schwann cells produced a Schwann cell substrate that aligned along explant axons and formed a monolayer around the explants (Figure 3A). After a single day of triple co-culture, the mESC-derived motor neurons appeared to contact explant axons (Figure 3B).

**Figure 3:**
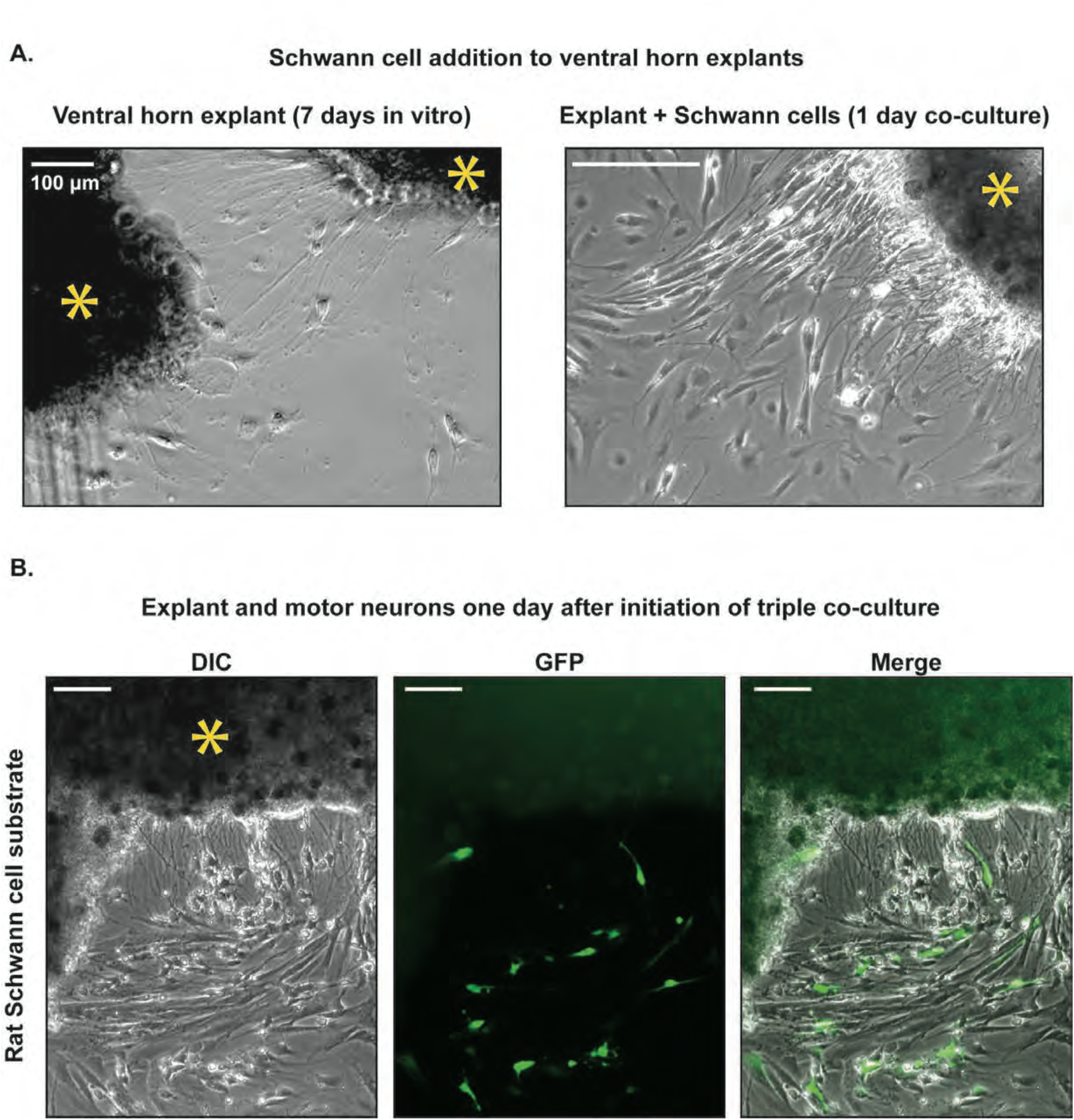
Rat spinal cord explant to differentiated motor neuron synaptogenesis A. Visualization of Schwann cell addition to explants. Axons are seen extending from the spinal cord explant (the same explant is visible in the upper left of each image) after one week *in vitro* (left). One day after addition of primary rat Schwann cells, the Schwann cells are see aligning with explant axons and creating a monolayer substrate (Schwann cells are identifiable by characteristic morphology). B. Live GFP and DIC images of the explant/glia/derived-motor neuron co-cultures (20x) one day after the initiation of triple co-culture. (*) indicates explant. Scale bars represent 100 μm.

To test electrochemical connectivity between rat spinal cord and mouse derived motor neurons, the triple culture was matured for seven days, loaded with Fura2, and arranged for electrical stimulation (Figure 4A). By combining all three of these components, it was hypothesized that *in vitro* studies of synaptogenesis could inform as to the possibility of rat lower motor neuron to mouse derived motor neuron synaptogenesis. If the ventral horn explant and motor neurons were synaptically connected, then electrical stimulation of the explant should lead to neurotransmitter mediated depolarization of the motor neurons. To avoid the possibility of electrical depolarization of the explant also causing electrical depolarization of motor neurons (rather than chemical depolarization), cells were stimulated in the presence and absence of glutamatergic and cholinergic synaptic blockers whereby the depolarization of chemical synapses would be expected to be abrogated by blockade.

**Figure 4:**
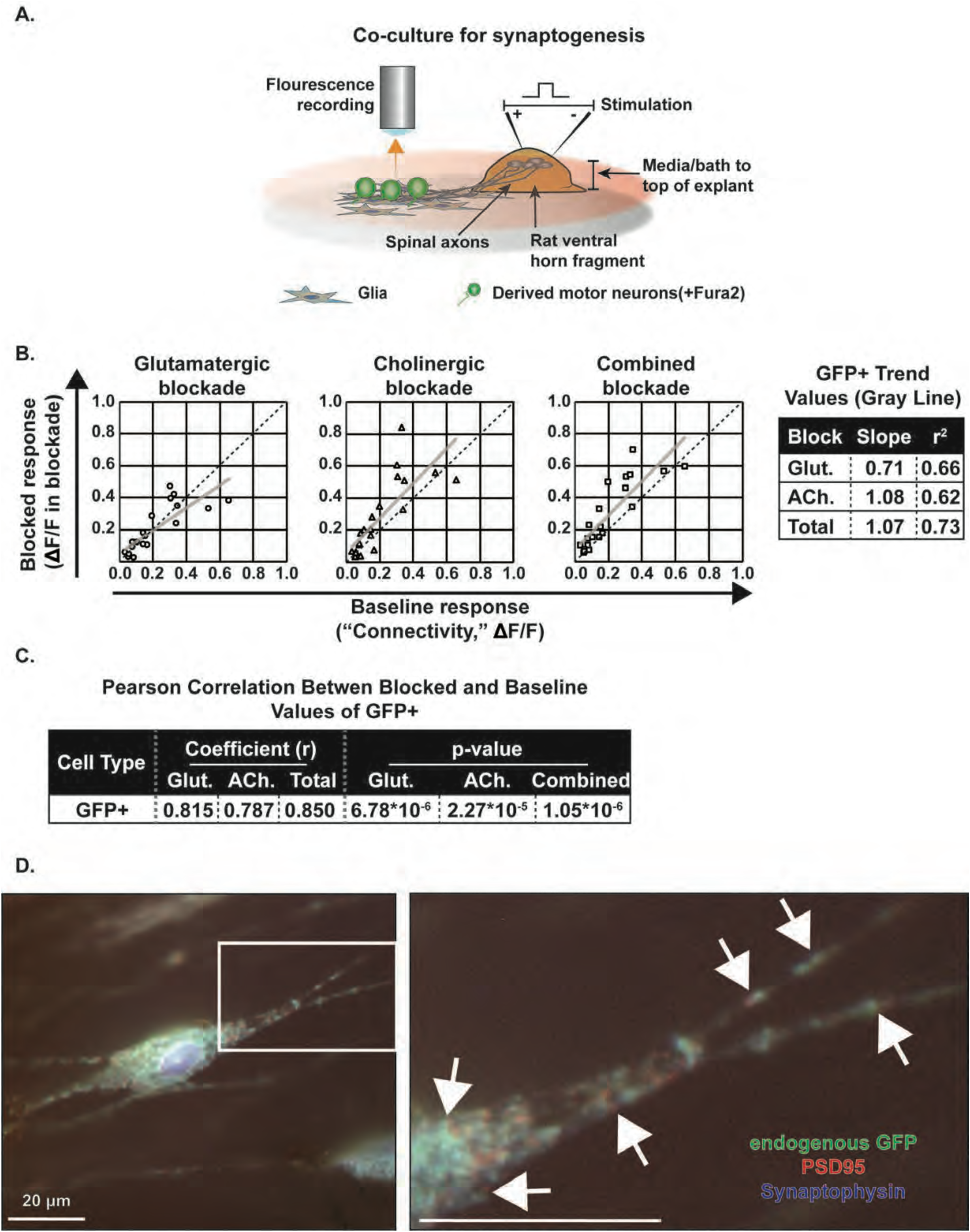
Rat spinal cord explant to differentiated motor neuron synaptogenesis (**A**). Schematic of calcium imaging set up demonstrating triple culture of ventral horn explant, Schwann cells, and derived motor neurons. The explant was electrically stimulated while visualizing for fluorescence of the calcium sensitive dye. Stimulation was performed in baths of artificial cerebrospinal fluid with and without glutamatergic and cholinergic blockade (**B**). Correlation diagrams of all GFP^+^ cells in each blockade shows a negative correlation with glutamatergic blockade, but positive correlation with cholinergic and combined blockade. Line of best fit shown in gray with parameters reported at right **(C)**. Pearson’s correlation for GFP^+^ cells in each blockade condition. N= 21 (GFP^+^), Bonferroni correction for multiple comparisons defines significance as P<0.0083. (D). After electrical stimulation and synaptic blockade studies, the cells were stained for markers of glutamategic synapses: synaptophysin (blue) and PSD95 (red). Some, but not all, derived motor neurons (endogenous GFP reporter, green) displayed staining consistent with synaptic structures adjacent red/blue staining within green cells). Boxed regions highlight expanded region of neurites at right. Images are of a single plane in a Z-stack of the cells, acquired at 63x. Scale bars represent 20 μm.

To assay synaptic connectivity between rat spinal cord explant sections and mESC-derived motor neurons, the explant section was electrically stimulated (bipolar 18V, 36 mA, 100 μS pulse at 18 Hz for 10 sec) while monitoring the Fura2-AM loaded, mESC-derived (GFP^+^) motor neurons for calcium transients with and without reversible glutamate (NBQX and CPP), acetylcholine (mecamylamine), or combined (NBQX, CPP, and mecamylamine) receptor antagonists.

When the electrically evoked depolarization in the blockade solution was plotted as a function of the initial, unblocked depolarization, each cell’s synaptic connectivity was assayed: if a cell was synaptically connected, its blocked depolarization should be less than its baseline. A stronger baseline connection should have a more robust attenuation with blockade. Indeed, the blocked versus unblocked response of GFP^+^ cells with the glutamatergic, cholinergic, and combined blockade, demonstrated a negative correlation with glutamatergic blockade (slope less than one), but a positive correlation (slope more than one) with the cholinergic and combined blockade (Figure 4B). A Pearson’s correlation with a Bonferroni correction for multiple comparisons confirmed a statistically significant correlation (Figure 4C). These data suggest GFP^+^ cells are synaptically connected to the spinal cord explant in a glutamatergic dependent manner. Immunofluorescent staining of PSD95 and synaptophysin confirmed synaptic structures in some GFP^+^ cells (Figure 4D). The increased calcium response in GFP+ cells to electric stimulation of the spinal cord explant with cholinergic blockade is thought to be due to inhibition of inhibitory cholinergic interneurons.

### In vivo transplantation and facilitation of regeneration

The survival of transplanted, mESC-derived motor neurons and their integration with host machinery were tested with imaging and functional studies. Following denervation, a two- or three-month delay between transplant and repair was used for the experiments designed to test the feasibility of relay formation and effect of transplanted cells to provide support for endogenous cells, respectively (Figure 5A). Robust denervation of the tibial nerve was completed through quadruple ligation, transection, and stump retroflexion (Figure 5B). Immunosuppressed host tibial nerve stumps were then transplanted with 2*10^5^ cells divided at two sites 5 and 2mm distal to the end of the stump (Figure 5C). As a negative control, denervated animals were also transplanted with 2*10^5^ cells that had been boiled for fifteen minutes to produce heat-killed control samples that would be biologically inert while still maintaining their immunogenic potential in the host.

**Figure 5.**
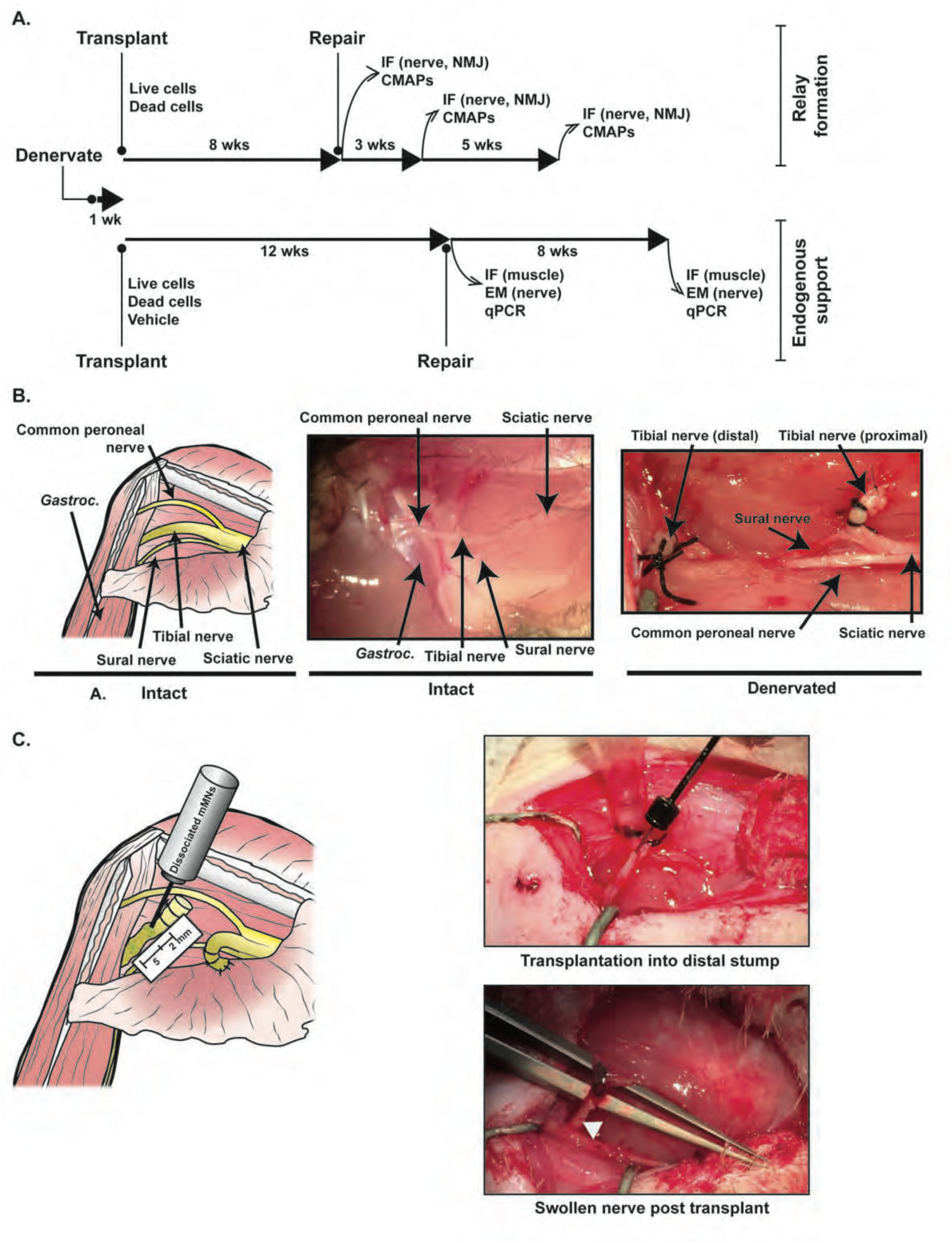
In vivo denervation and transplantation. (**A.**) Overview of in vivo experimental design to examine both the relay and endogenous support hypotheses. Two and three-month delay between transplant and repair intended to produce chronic denervation phenotype. IF: immunofluorescent staining; CMAPs: Compound muscle action potentials from intrinsic foot muscles; EM: Electron microscopy; qPCR: quantitative PCR of active Schwann cell markers. (**B.**) Schematic (left) and gross anatomy (middle) of the lateral approach to the sciatic nerve in the upper left hind limb of a rodent highlighting normal anatomy and the denervation protocol *in vivo* experiments. Sciatic notch is to the right in all images. Post-surgical anatomy illustrates denervation with quadruple ligatures around the tibial nerve bisected with full transection and both the proximal and distal stump sutured to adjacent muscles in opposing orientation. (**C.**) Schematic of cell injection into distal nerve stump (left) and gross images (right) of injection (top) into two sites in the denervated distal tibial nerve stump. After injection (bottom), the distal stump is visibly swollen (arrowhead).

Distal tibial nerve samples were stained for axons (β-III-tubulin) at five and eight weeks after transplantation (with no repair), where many axons were visible at the graft site and more distally in live, but not dead, cell transplanted animals suggesting that transplanted mESC-derived motor neurons were able to integrate into the host environment and extend axons (figure 6A). Eight weeks after transplantation, the endogenous GFP was no longer visible, but amplification by anti-GFP immunofluorescence demonstrated cells bodies at five and eight weeks after transplantation (Figure 6A, left column, and Figure 6B). While axons were identified to extend distally, examination of the neuromuscular junction of the soleus muscle, distal to the tibial nerve, demonstrated the presence of axons without clear neuromuscular junction formation (Figure 7A). Additionally, electrodiagnostic studies did not reliably demonstrate a compound muscle action potential following stimulation of the transplantation site eight weeks after transplantation (Figure 7B).

**Figure 6.**
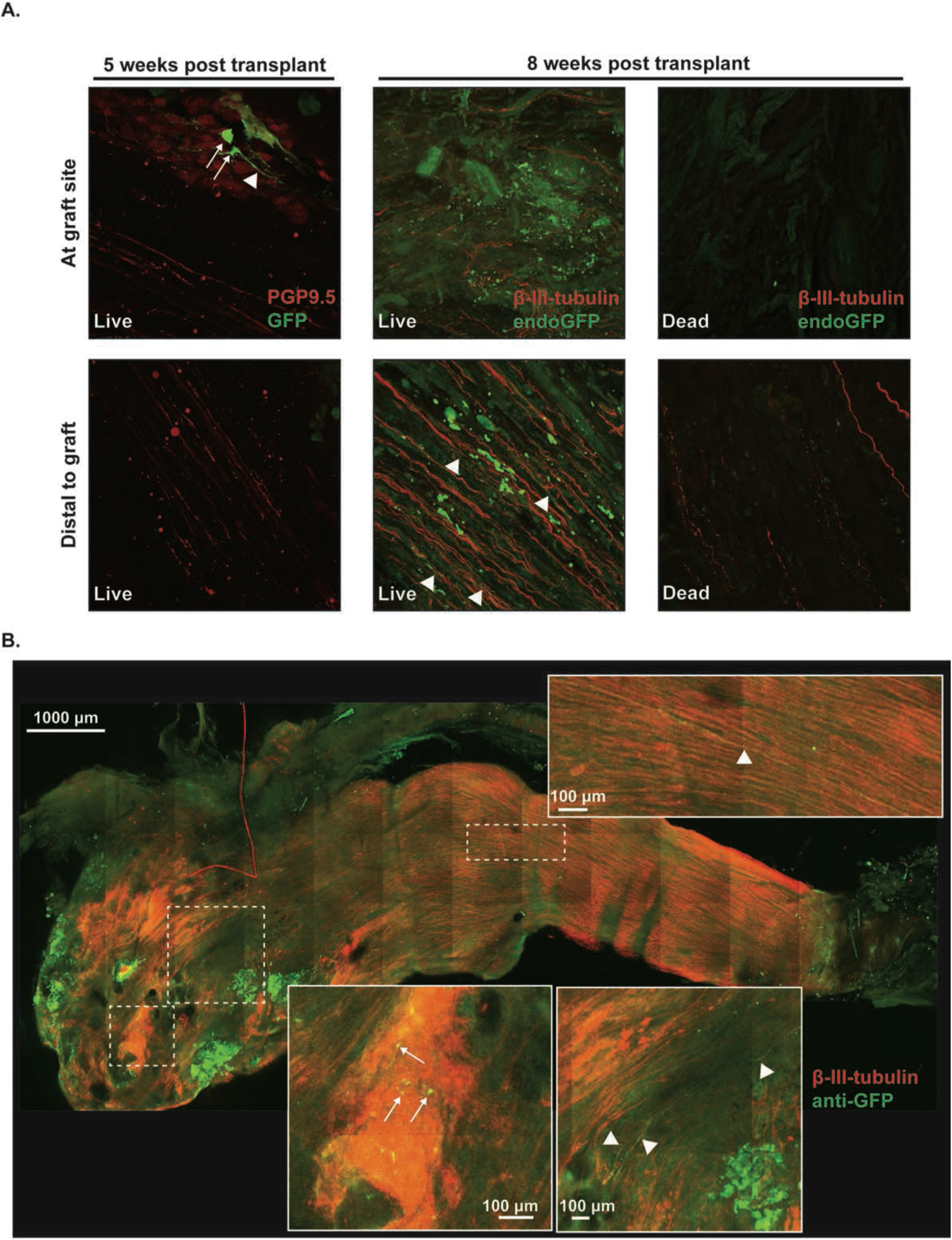
Engraftment of transplanted motor neurons into host tibial nerve. (**A.**) Confocal maximum intensity Z-projection images (40x and 20x as indicated) at and distal to the graft site of transplanted differentiated Hb9:GFP motor neurons in the denervated tibial nerve of immunosuppressed rat hosts five and eight weeks after transplant. Note that the Hb9:GFP reporter has been inactivated eight weeks after transplant. Red: anti-PGP9.5 (axonal stain); Green: α-GFP for the five-week time point. Red: anti-β-III-tubulin; Green: endogenous GFP reporter for the eight-week time point. (**B.**) Confocal Z-stack projection mosaic of 10x images from CUBIC cleared, transplanted nerve stump 8 weeks after live derived motor neuron transplantation, with staining against β-III-tubulin (red) and GFP (green) shows cell bodies (arrows) with distal projections (triangles). Dotted outline indicated enlarged regions in insets.

**Figure 7.**
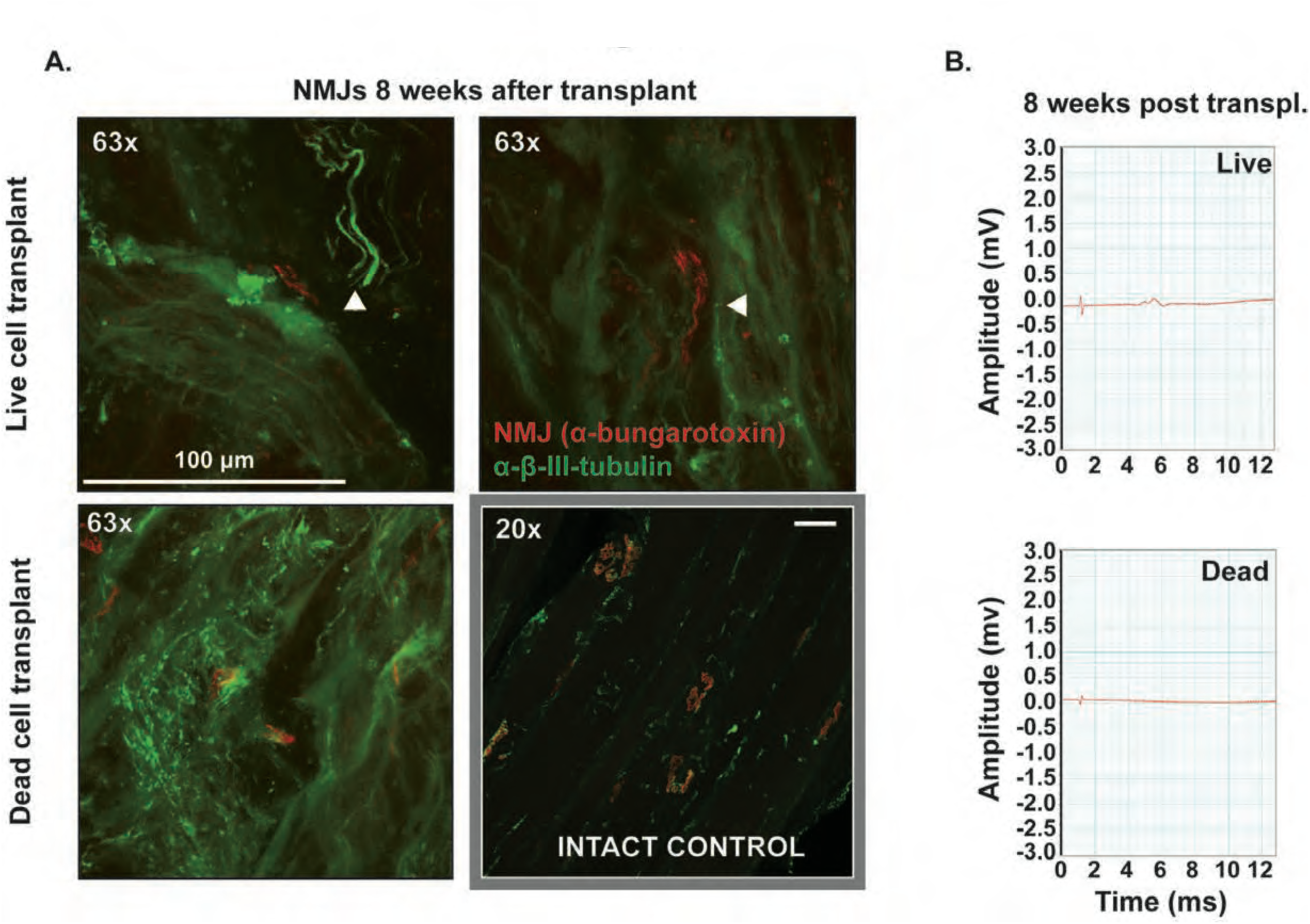
Distal axon projection without functional neuromuscular junction formation. (**A.**) Axons (anti β-III-tubulin; Green) from live (but not dead) transplanted cells are visualized at the neuromuscular junction (alpha-bungarotoxin; Red) in sections of the soleus muscle eight weeks after transplantation but prior to repair on 63x confocal Z-projection images. However, there is no evidence of neuromuscular junction innervation. A 20x confocal Z-projection of an intact muscle is shown to illustrate normal morphology. (**B.**) Nerve conduction studies of the transplanted nerve stump in live and dead cell transplanted nerves show no reliable compound muscle action potential evoked in the intrinsic foot muscles.

Following an eight- or twelve-week delay, the transplanted nerve stump was sutured to a freshly transected peroneal nerve. Axonal staining demonstrated regeneration of host axons into the distal stump, but without clear entrance into the transplant site (Figure 8A). To determine the effect of transplanted cells on host regeneration, intrinsic foot muscle CMAPs were obtained from direct sciatic nerve stimulation 3-5 mm proximal to the common peroneal-tibial nerve repair site three and eight weeks after repair (11-16 weeks after transplant). At eight weeks following repair, these CMAPs were robust in live but not dead cell transplanted animals, and an increase in the amplitude was demonstrated over time, suggesting the mESC-derived motor neurons facilitated regeneration after a two-month delay between transplantation and repair (Figure 8B). Quantification of the peak-trough amplitude of the intrinsic foot muscle CMAPs after both a two- and three-month delay between transplant and repair confirmed the initial observations (Figure 8C). Indeed, a statistically significant increase in amplitude in live animals from zero to 56 days post-repair was observed in both the two and three-month delay, and the amplitude 56 days post-repair was significantly greater in live versus dead cell transplant groups. Staining for synaptic markers at the graft site was negative and there was no reliable electrodiagnostic evidence of relay formation (ie CMAP duration prolongation or second discharges), which prompted investigation into the effect of the transplanted cells on the host Schwann cell phenotype.

**Figure 8.**
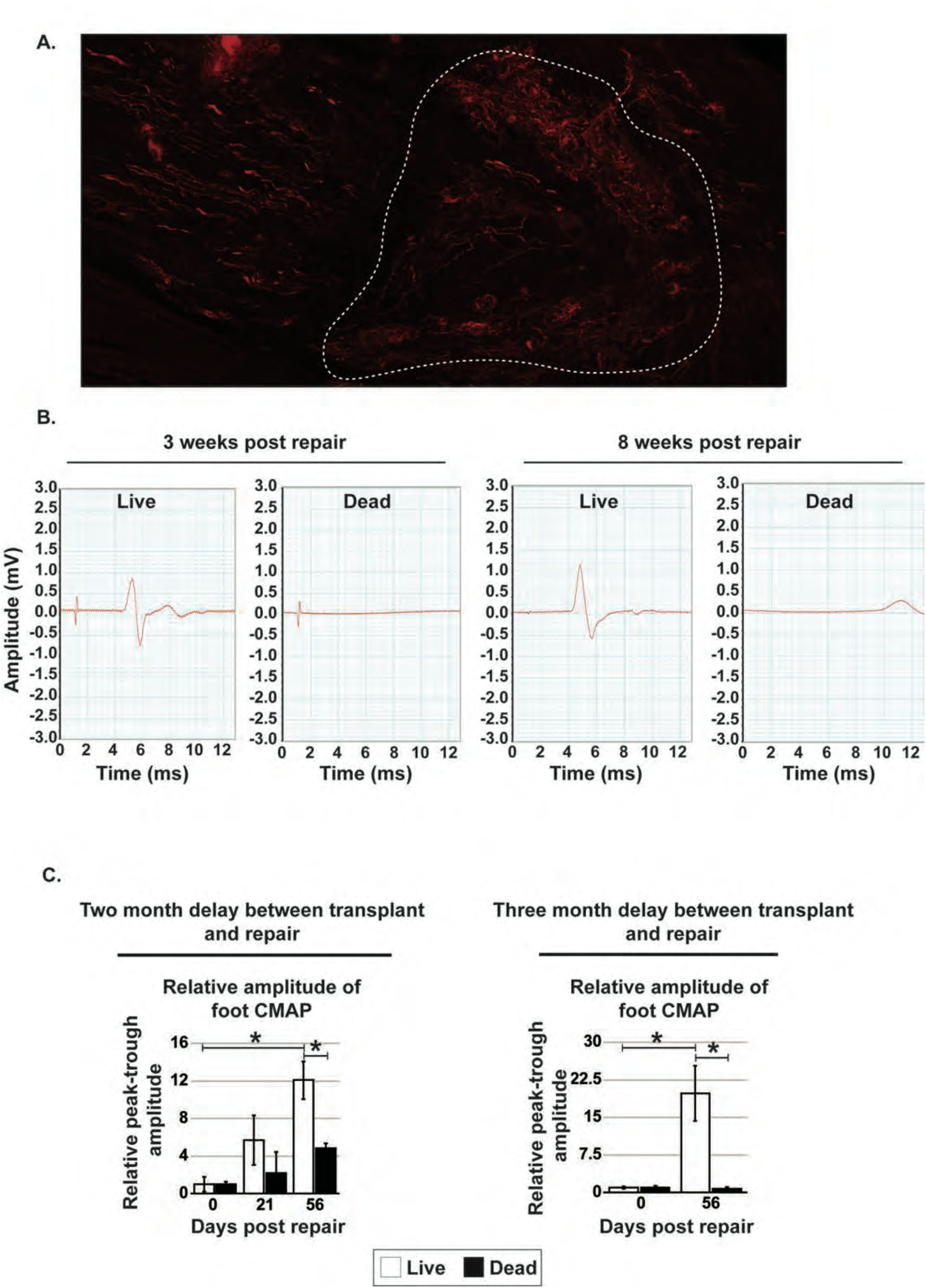
Cell transplantation increases regeneration of a chronically denervated nerve (**A.**) Confocal max intensity Z-projection of β-III-tubulin staining at the graft site in a longitudinal section of the tibial nerve eight weeks after repair (sixteen weeks after transplant). Proximal nerve is to the left, white dotted line denotes transplant site. (**B**). CMAP recordings from the intrinsic foot muscle with direct stimulation of the sciatic nerve of a rat three weeks and eight weeks following repair of the transplanted tibial nerve. Eight weeks prior to the repair, the animal had been transplanted with live or dead differentiated motor neurons. (**A**). Quantitation of intrinsic foot CMAP peak-trough amplitude relative to values at the time of the repair. Left panels were from animals with a two-month delay between transplant and repair, while right panels were from animals with a three-month delay. Error bars are SEM. *=p<0.05 by ANOVA with Tukey’s post-hoc analysis, α=0.05 for comparisons of a single treatment group across all three time points in two-month delay animals; For three month delay, Student’s two-tailed t-test with equal and unequal variance determined by f-test for comparison between treatment groups at a given time point. n=5, 5, 4 animals for live cell and n=2,3,3, animals for dead transplant cell animals at 0, 21, and 56 days after repair in two-month delayed animals, respectively. n=6, 6, animals for live cell and n=7,8 animals for dead cell transplants with three-month delay between transplant and repair.

### Schwann cell support of regeneration

The ability of transplanted live, mouse embryonic stem cell (mESC)-derived motor neurons to maintain the capability of Schwann cells to adopt an active, pro-regenerative phenotype following repair was examined by imaging and gene expression analysis in live cell, dead cell, and vehicle transplanted denervated nerves. Since these studies were intended to examine the effects of the transplanted cells on host Schwann cells, instead of host axons, a longer delay (twelve weeks instead of eight weeks) was used between transplant and repair, to ensure complete adoption of the chronically denervated phenotype, as discussed previously (Figure 5A). Electron micrographs of tibial nerve cross sections twelve weeks after transplant showed Schwann cells surrounded by collagen and fibroblasts in all three transplant groups (Figure 9A). Unmyelinated axons were also readily identified. Not pictured, but present in other sections, were myelinated fibers. Quantitation of axons (myelinated and unmyelinated), Schwann cells, and fibroblasts in the electron micrographs was performed both at the time of repair and eight weeks after repair (12- and 20-weeks following transplant, respectively). Surprisingly, in the distal nerve (where the EM sections were obtained), there were no significantly different cell counts between treatment type or time points (Figure 9B), although live cell transplants tended to have more axons and Schwann cells and fewer fibroblasts at the time of repair. While the cell counts did not differ, the existing Schwann cells may have been phenotypically different, which necessitated an examination of gene expression.

**Figure 9:**
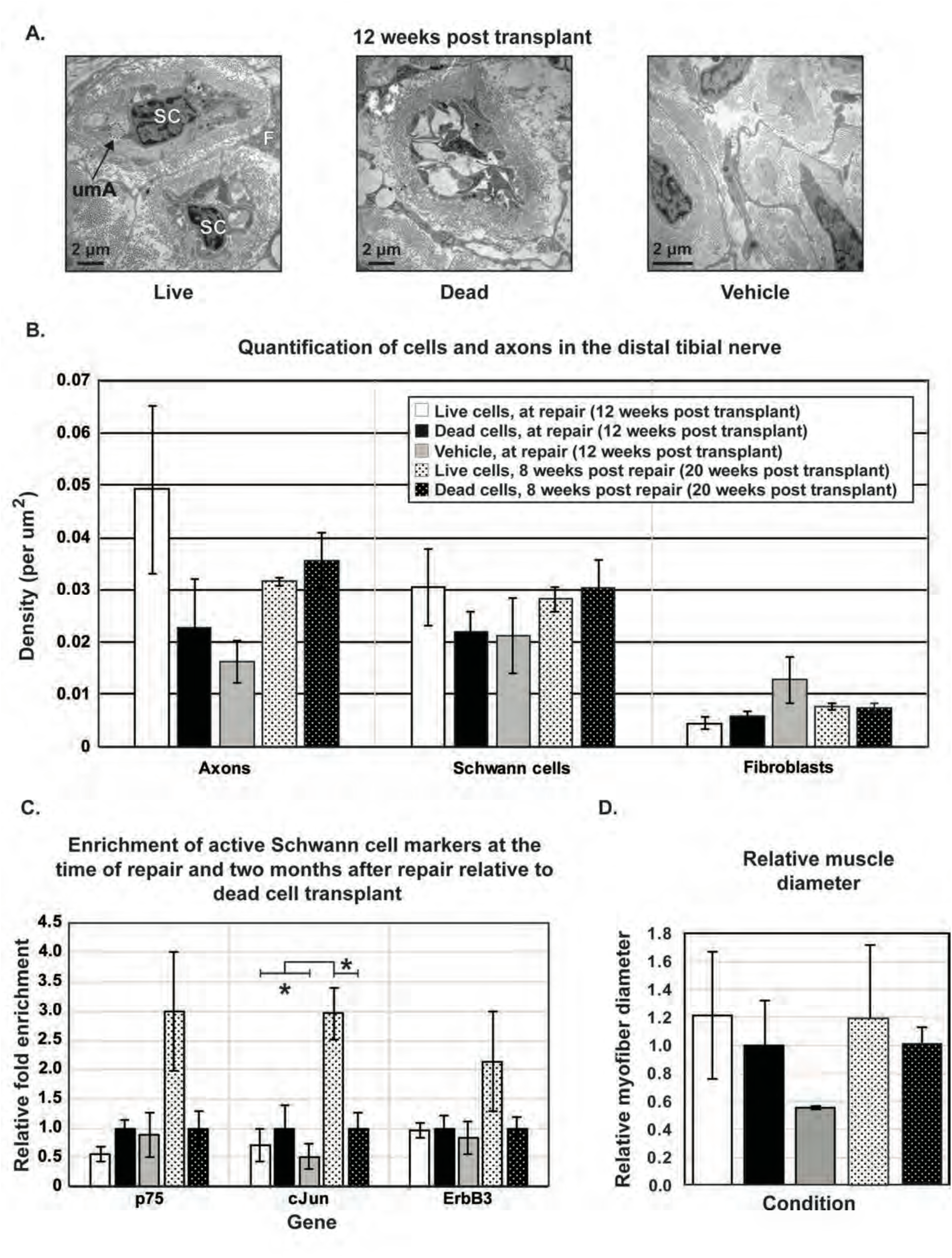
Transplanted cells maintain host Schwann cell activation potential to facilitate regeneration (**A**). Electron micrographs (8000x) of tibial nerve cross sections in all three groups at the time of repair, three months after transplant with Schwann cells (SCs), fibroblasts (F), and unmyelinated axons (umA, outlined) visible. (**B**). Quantification of axons (myelinated and unmyelinated), Schwann cells, and fibroblasts in electron micrographs of tibial nerve cross sections at the time of repair and eight weeks after repair. No statistically significant difference between treatment groups and time points by ANOVA, α=0.05. n=2 and 4 animals for live and dead cell transplants, respectively, for both time points. n=4 animals for vehicle control. (**C**). Fold enrichment of the same genes in tibial and (repaired) sciatic nerve samples in live and dead cell transplant groups at the time or repair and eight weeks after repair, following a three-month delay between transplant and repair. Fold enrichment is relative to intact side, relative to the dead cell transplant at each time point. *=p<0.05 for comparison across groups within a single gene by ANOVA and Tukey’s post-hoc analysis, α=0.05. n=4, 3 animals for live and dead cell transplants, respectively, at both time points. n=2 animals for vehicle transplant. (**D**). Quantitation of myofiber diameter relative to intact diameter at the time of repair and eight weeks after repair. Values are relative to dead cell transplant at each time point. No significant difference by ANOVA, α=0.05, n=3, 4 animals for live and dead cell transplant groups, respectively, at both time points. n=3 for vehicle transplant animals. Error bars are SEM.

To this end, the Schwann cells in the live cell, dead cell, and vehicle transplanted nerve were examined following a three-month delay between transplant and repair. Three proteins, p75, ErbB3, and c-Jun are known to increase in expression during Schwann cell activation and were used to characterize the state of the host Schwann cells (Carroll et al., 1997; Müller and Stoll, 1998; Arthur-Farraj et al., 2012). When the expression of these genes was assayed in the transplanted tibial nerves at the time of repair and two months after repair, all three genes were enriched in live cell transplanted animals only, two months after repair (Figure 9C). While all three genes were enriched, c-Jun was statistically significantly elevated at eight weeks following repair versus at the time of repair.

In addition to an effect on Schwann cells, the transplanted population could have facilitated regeneration through modulation of distal musculature. To evaluate muscle atrophy following the twelve-week delay between transplant and repair, myofiber diameter was measured in the soleus muscle from transplanted animals at the time of repair and two months after repair. There was no significant difference in myofiber diameter, although there a trend of vehicle tending to be lower than the live or dead cell transplants (Figure 9D). These data suggest the transplant facilitated maintenance of regenerative capacity was not due to an effect on muscle.

## Discussion

Since chronic denervation is due to loss of Schwann cells from extended denervation and subsequent atrophy, transplantation of neurons into a denervated nerve was predicted to slow the conversion to a chronic denervation phenotype through relay formation across the graft as well as through support of the endogenous pro-regenerative Schwann cells. The former mechanism could reduce time of regeneration drastically, as the regenerative distance would be covered from multiple points that would then be connected, while the latter would reinforce Schwann cells against degeneration.

To this end, motor neurons were derived from mouse embryonic stem cells, injected into the distal stump of a transected nerve then nerve repair delayed at two or three months for chronic denervation followed by a two-month regeneration window following repair. The differentiation resulted in a heterogenous mix of motor neurons, non-motor neurons, neural stem cells, and glia, which was considered ideal for transplant for glial support of the neurons. The motor neurons in this population were found to express machinery for both glutamatergic (vGLUT, PSD95) and cholinergic (ChAT) synapses. Co-culture of these neurons with a rat spinal cord explant demonstrated rat spinal cord to mouse motor neuron synaptogenesis in a glutamatergic dependent manner. Furthermore, some of these derived motor neurons elaborated appropriate synaptic machinery on staining. All these data suggest derived motor neurons can serve a post-synaptic target for regenerating host fibers, a necessary substrate for relay formation from spinal cord to muscle. Importantly, neither glutamatergic kainate nor metabotropic receptors were blocked during neurotransmitter inhibition, which may explain the failure to completely abolish calcium transients (Tölle et al., 1993). Nonetheless, the significant reduction in depolarization following inhibition of glutamatergic transmission suggested some functional connectivity between explant and mESC-derived motor neurons. The presence of ChAT positive but GFP negative neurons was consistent with presence of inhibitory interneurons, which was confirmed on electrical stimulation whereby cholinergic blockade increased depolarization. The data did not definitively demonstrate direct rat motor neuron to mouse motor neuron connectivity, as interneurons could still be critical in relay formation. Indeed, the variable response of cells to single versus combined blockade suggest multiple cells and their respective synaptic identifies are likely involved in the explant to motor neuron connectivity. Such distinctions may only be studied by direct cell-to-cell electrophysiological studies. Since this distinction was not critical for the formation of a relay, per se, it was considered to be outside of the scope of this study.

After successful demonstration of derived motor neuron and rat ventral horn synapse formation *in vitro*, *in vivo* experimentation was completed to determine if intraneural motor neuron transplantation could maintain regenerative capacity in a chronically denervated nerve. Mechanistic studies were completed through two different methods. To study relay formation, the unsorted, differentiated motor neuron population was injected into the distal nerve stump of a transected tibial nerve and allowed to engraft for two months. At that time, the peroneal nerve was transected and the proximal stump of the peroneal nerve sutured to the previously injected distal stump of the tibial nerve, thereby permitting acute regeneration through a chronically denervated nerve segment. Regeneration occurred for two months, with electrodiagnostic testing at the time of repair and two months after repair. The ability of the transplanted cells to support endogenous Schwann cells was with an identical protocol, except repair was delayed for three instead of two months, and nerve morphometry and gene expression data performed at eight weeks post repair.

Successful relay formation *in vivo* necessitates grafted cells to synapse onto muscle as well as to serve as postsynaptic targets to regenerating host axons. While the latter was demonstrated *in vitro* (Deshpande et al., 2006), transplanted cells axons were not observed to innervate neuromuscular junctions in our studies, and there was no reliable electrodiagnostic support for such. Eight weeks following nerve repair, host axons were seen to enter the chronically denervated stump, but not to clearly innervate the cell graft, with no evidence of synaptogenesis. This difference from the in vitro model may be due to fibrosis or scarring at the transplant site and/or the relatively decreased density of derive motor neurons in the 3D nerve space versus the culture plate. Nonetheless, serial CMAPs of the intrinsic foot muscles with stimulation above the repair site showed robust regeneration following nerve repair in live but not dead cell transplanted animals after two and three months of denervation prior to repair, demonstrating that transplantation of mESC-derived motor neurons maintains the regenerative capacity of chronically denervated nerves in the absence of relay formation.

To evaluate if the transplanted cells may support host pro-regenerative cells, the distal most aspect of the distal stump of the transplanted, denervated nerve was imaged by electron microscopy at the time of repair (three months after transplant of dead cells, live cells, or vehicle) and two months after repair. This imaging showed no difference in axon or cell number between treatment group at either time point, likely a function of sample size, since the live cell transplants were limited by side effects in these animals. The axons observed in the vehicle transplanted animals may have been from the vasa nervorum nerve plexus sprouting which has been shown to still be present after three months of denervation (Hoke et al., 2001). Notably, the difference between live and dead cells became more similar eight weeks after repair, despite the previous observation that regeneration is much more robust in live cells than dead cell transplants. Interestingly, a recent study of peripheral nerve regeneration following transection with repair at different delays found myelin thickness and diameter did not accurately represent recovery (Ikeda and Oka, 2012), (Ikeda and Oka, 2012), so the failure to detect many large, myelinated fibers in the live cell transplant group does not necessarily conflict with the observed improvement in regeneration. Indeed, a numeric difference may not be necessary to facilitate regeneration, whereby a phenotypic difference of host cells may be sufficient. Additionally, it is possible that the graft has a more robust effect on the number of host cells more proximal to the graft site than the sections examined here. To examine the phenotype of the host Schwann cells (regenerative versus non-regenerative) Schwann cell gene expression was studied.

While the Schwann cells express similar levels of p75, c-Jun, and ErbB3 gene transcripts at the time of repair, after the new insult of repair, the Schwann cells of live, but not dead cell or vehicle, transplanted animals were capable of adopting an active phenotype with the concomitant increase in transcript levels of the three genes. Thus, while at baseline (three months after transplant) live and dead cell transplanted nerves appeared to contain similar densities of Schwann cells, as was observed in the electron microscopy, Schwann cells were better preserved by the live cells, suggesting live cell transplants supported host cells.

In addition to Schwann cell inactivation and loss, muscle atrophy has been accepted as one of the factors limiting peripheral nerve regeneration after chronic denervation (Gordon et al., 2011). There was no significant difference in myofiber diameter between live, dead, and vehicle treatments, although there were trends to suggest both live and dead cells may provide trophic support for muscle since vehicle transplants demonstrated more atrophy, but, given the lack of statistical significance, additional testing is necessary. Interestingly, myofiber diameter did not increase following repair. These data were consistent with the previously work showing transplanted neurons cannot restore atrophied muscle to a normal size (Kubo et al., 2009). While Kubo and colleagues suggested resistance to atrophy is effected through transplanted neuron innervation of NMJs, dead cell transplants were not examined, which is a particularly important control since the innate immune system has been demonstrated as a major facilitator of muscle regeneration (Chazaud, 2016). (Chazaud, 2016). Thus, the innate immune response to both live and dead cell transplants may have helped to maintain the muscle mass over vehicle, but additional testing is critical to determine if the observed trend is a true, statistically significant relationship, and to define the particular population of immune cells that may affect this response.

Finally, it should be noted that tumor formation was a common occurrence in all animals with live cell transplants, thought likely to be due to the presence of neural stem cells in the transplant population. This finding supports other work in cell-based therapies that recommends treatment with mitotic inhibitors (Magown et al., 2016).

Thus, intraneural transplantation of a mixed motor neuron population promotes regeneration in the chronically denervated nerve. While derived mouse motor neurons may serve as postsynaptic targets of rat lower motor neurons *in vitro*, robust synaptogenesis by regeneration fibers onto the transplanted cells does not occur. Instead, the maintenance of regenerative capacity in a chronically denervated nerve is due to maintenance of the endogenous Schwann cell population in a pro-regenerative state despite no statistically significant difference in Schwann cell number. Future work will focus on elucidate of the cells and molecules in the transplant that effect this benefit.

## Contributions

CRC and AH prepared the manuscript and figures. CRC and RM performed all experiments. CRC and AH designed all experiments. All authors reviewed the manuscript

## Acknowledgements

The authors would like to thank Qiang Shi and Brian Lin for their technical assistance during these experiments. The HBG3 Hb9:GFP mouse embryonic stem cells and mouse astrocytes were a kind gift from Dr. Nicholas Maragakis, Johns Hopkins School of Medicine (which were a gift from Dr. Thomas Jessell). Mouse astrocytes were prepared by the laboratory of Jeffrey Rothstein, Johns Hopkins School of Medicine.

## Funding

This work was supported by TEDCO 2010-MSCRFE-0194, 2014-MSCRFI-0715, and Dr. Miriam and Sheldon G. Adelson Medical Research Foundation.

## Conflict of Interest

None

## Notes

### Competing Interest Statement

The authors have declared no competing interest.

